# Cross-scale effect of microbiota dynamics on resistance

**DOI:** 10.64898/2025.12.04.692100

**Authors:** Lisa Pagani, Silvana Gloor, Silvio D. Brugger, Sebastian Bonhoeffer, Roger D. Kouyos

**Author notes:** These authors contributed equally to this work (Sebastian Bonhoeffer, Roger D. Kouyos).

## Abstract

Antimicrobial resistance (AMR) is often carried asymptomatically in the human gut microbiota, yet the role of this carriage in AMR persistence and transmission remains poorly understood. To address this gap, we developed a cross-scale model that links within-host microbiota dynamics of extended-spectrum beta-lactamase (ESBL)-producing and non-ESBL-producing bacteria with transmission across hospital and community settings. The model incorporates plasmid exchange, antibiotic effects, and host heterogeneity. It integrates diverse data sources, including microbiota dynamics, community antibiotic consumption, and detailed surveillance data from a large university hospital, complemented by literature estimates. This model identifies mechanisms that enable coexistence of resistant and sensitive strains at both individual and population levels, revealing host heterogeneity as a critical driver of resistance persistence. Unlike classic epidemiological models, our analysis finds a continuum of colonization states within individuals rather than discrete resistant or sensitive categories, and highlights that realistic detection thresholds may substantially underestimate resistance prevalence. Finally, we use this framework to evaluate interventions such as hospital screening and isolation, showing that their impact strongly depends on local transmission dynamics and resistance prevalence. By bridging within-host and population processes, our model provides a novel tool for investigating AMR spread and optimizing prevention strategies in healthcare and community environments.

## Introduction

Antimicrobial resistance (AMR) is a growing global health crisis. In 2021, AMR was associated with an estimated 4.71 million deaths [1]. AMR contributes not only to increased mortality but also to greater rates of treatment failure, prolonged hospital stays, and substantial economic burdens on healthcare systems globally [2, 3]. AMR is widely distributed across diverse environments, including hospital effluents, wastewater treatment plants, agricultural soils, and natural aquatic systems [4–6]. Beyond the environment, AMR is carried by animals and humans, where it can persist within their microbiota [7]. The human microbiota constitutes a rich and diverse ecosystem composed of bacteria, fungi, viruses, and archaea, structured into site-specific communities such as those of the gut, skin, oral cavity, and respiratory tract [8]. Among these, the gut microbiota is the largest and most densely populated microbial community of the body, harboring up to 10^14^ microorganisms. [9–11]. Genomic approaches demonstrate that the relative abundance and diversity of AMR are particularly high in the gut, underscoring its role as a major reservoir of resistance [12].

The gut microbiome provides an ideal environment for the emergence, spread, and persistence of resistance [13]. It may be repeatedly exposed to bystander selection pressures during antimicrobial treatment targeting specific pathogenic infections, resulting in collateral effects on commensal communities. The gut harbors high microbial diversity and density, and it facilitates horizontal gene transfer (HGT), including plasmid exchange of resistance genes [14, 15]. Resistant microorganisms can persist asymptomatically within the gut, evade detection, and be transmitted to others via direct or environmental routes. Although frequently asymptomatic, these microorganisms can translocate to the bloodstream or internal organs, or disseminate to other body sites, causing infections with resistant organisms for which first-line therapies are ineffective [16–19].

This dynamic is particularly concerning in the case of Gram-negative bacteria producing extended-spectrum beta-lactamases (ESBLs), which are enzymes that confer resistance to most beta-lactam antibiotics and are commonly carried on mobile genetic elements such as plasmids and transposons [20, 21]. Because of this localization, the acquisition of ESBL genes is predominantly mediated by HGT, enabling dissemination within and between bacterial species [22]. Key ESBL-producing species include *Escherichia coli, Klebsiella pneumoniae, Pseudomonas aeruginosa*, and *Enterobacter* species, which are frequent causes of healthcare-associated infections and increasingly difficult to treat [21]. These species are gut commensal bacteria and estimates suggest that over 1.5 billion people are colonized [23]. Understanding the dynamics of commensal ESBL carriage is therefore essential, as it represents a reservoir from which resistant infections originate and through which resistance can disseminate across hosts and environments [24].

Despite increasing recognition of the gut microbiome’s role in AMR, the integration of within-host microbial dynamics into AMR epidemiological models remains limited. Numerous models have investigated AMR transmission in healthcare and community settings [25] or examined ecological selection of AMR within the microbiome [26–33]. However, only few have explicitly connected these scales to understand how within-host dynamics influence epidemiological patterns. [34]. Incorporating mathematical modeling into the study of microbiome resistance offers unique opportunities: models provide a mechanistic framework to formalize biological hypotheses, capture complex interactions, and predict system behavior under diverse conditions. This approach allows simulation of scenarios that are experimentally challenging and can generate testable predictions, thereby guiding empirical work and optimizing the allocation of research efforts.

In this study, we develop a novel cross-scale model that links within-host gut microbiota dynamics, including antibiotic exposure, plasmid resistance, and HGT, with epidemiological transmission in both hospital and community settings. Applying this framework to ESBL-producing Gram-negative bacteria, we show that population heterogeneity drives persistence of resistance, and that detection thresholds constrain our ability to accurately identify ESBL carriers. We further demonstrate that interventions such as hospital isolation, and gut decolonization can mitigate but not fully prevent resistance dissemination. Our work bridges the gap between ecological and epidemiological perspectives, providing new insight into resistance persistence, dissemination, and the impact of intervention strategies.

## Results

### Model

We develop a cross-scale computational model of the gut microbiota to characterize the dynamics of commensal bacteria and the dissemination of antibiotic resistance plasmids. Our model builds on the framework established by Tepekule *et al*. [28], which simulates gut microbiota dynamics with a focus on plasmid resistance. In contrast to the original formulation, which incorporates four major bacterial phyla, we tailor our approach to specifically model the Proteobacteria phylum (Fig. 1). This refinement is motivated by the high prevalence of antibiotic resistant Proteobacteria, making it a critical group for investigating resistance transmission dynamics [35]. This simplification of the microbiota dimension enables us to capture the essential within-host processes while making the model tractable for extension to the population scale.

**Figure 1.**
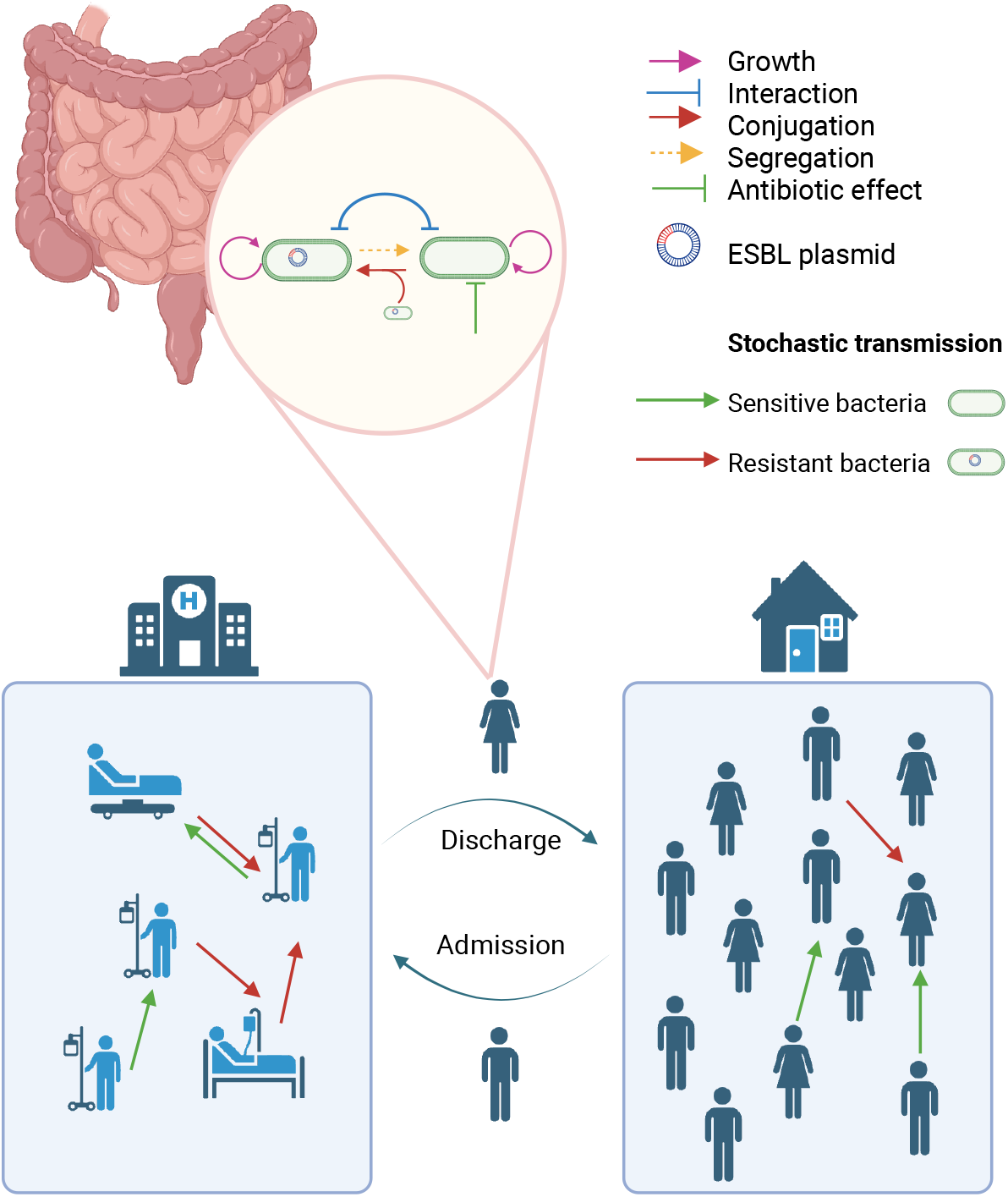
Overview of model dynamics. Schematic summary of the hybrid model combining within-host microbiota dynamics and between-host transmission processes across hospital and community settings. The upper part illustrates the within-host dynamics of sensitive (S) and resistant (R) Proteobacteria, while the lower part represents transmission routes and movements in both environments. Created in BioRender.

In our formulation, individuals carry gut populations of non-ESBL-producing (S) and ESBL-producing (R) Proteobacteria, and we refer to these as *sensitive* and *resistant* bacteria throughout the manuscript. These populations evolve through growth, competition, plasmid conjugation, plasmid loss, and antibiotic selection (Fig. 1). At equilibrium, the total number of gut sensitive bacteria is approximately 3 10^11^. The bacterial dynamics are embedded within a structured host population comprising two fixed-size environments: hospital and community (Fig. 1). This structure allows us to account for different transmission patterns, antibiotic exposure rates, and admission/discharge flows.

To capture realistic features of antibiotic resistance dynamics, we incorporate two critical components. First, the model includes heterogeneity in community antibiotic consumption, based on empirical data showing that a small fraction of individuals accounts for the majority of outpatient antibiotic use [36]. Second, we model non-random hospital admission, where individuals with higher outpatient antibiotic consumption are more likely to be hospitalized, reflecting increased fragility or infection risk [37]. The state incorporating community consumption variability and non-random transitions is referred to as the heterogeneous model, while the absence of such variability defines the homogeneous model.

The model operates with deterministic state updates of the gut microbiota occurring daily, while incorporating stochastic transmission events within each environment. During each transmission event, a fixed inoculum size of 10^4^ bacterial cells are transferred from the donor to a randomly chosen recipient within the same environment (Fig. 1). This value is based on empirical observations indicating that transmission events typically involve bacterial loads on the order of thousands of cells [38]. The composition of the transferred bacteria, either all sensitive or all resistant, is determined by the donor’s gut bacterial composition, calculated as 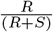, where *R* and *S* represent the donor’s resistant and sensitive bacterial populations. It should be noted that this model abstracts the transmission process without specifying the exact mechanisms involved, acknowledging that transmission may occur via both direct and indirect pathways.

We parameterize the model to represent ESBL-producing Proteobacteria as a clinically relevant case study. ESBL-producing bacteria are considered critical priority pathogens by the World Health Organization due to their association with multidrug resistance and limited treatment options [21]. To realistically capture ESBL dynamics, we further parameterize the model using a previously published gut microbiota framework [28], a detailed surveillance dataset from the University Hospital Zurich, and outpatient antibiotic consumption data from the Swiss Center for Antibiotic Resistance [39]. In our framework, exposure to beta-lactam antibiotic selects for plasmid-carrying ESBL bacteria [40]. For simplicity, the model does not include the consumption of other antibiotic classes that could co-select for both sensitive and resistant bacteria.

Together, this model provides a mechanistic framework to investigate how within-host plasmid dynamics scale to population-level resistance spread under realistic antibiotic use patterns. Full implementation details and governing equations are provided in *Materials and Methods*.

### Resistance persists under population heterogeneity

The temporal dynamics of resistant colonization are examined across both hospital and community settings. We define an individual as colonized when at least one resistant bacterium is present in their gut. In the presence of population heterogeneity, resistance persists over time and ultimately stabilizes at high prevalence levels, reaching a plateau of approximately 99% colonization in the hospital and 81% in the community (Fig. 2). Notably, the probability of transmission in each setting plays a critical role in determining the overall colonization level by the end of the simulation (Fig. S1). However, resistance persists across all scenarios involving non-zero transmission rates. Only in the complete absence of transmission in both settings does resistance go extinct. In contrast, a model lacking population heterogeneity (no distinct categories for outpatient antibiotic consumption and hospital admission rates) fails to sustain resistance, underscoring the critical role of heterogeneity in resistance dynamics (Fig. 2).

**Figure 2.**
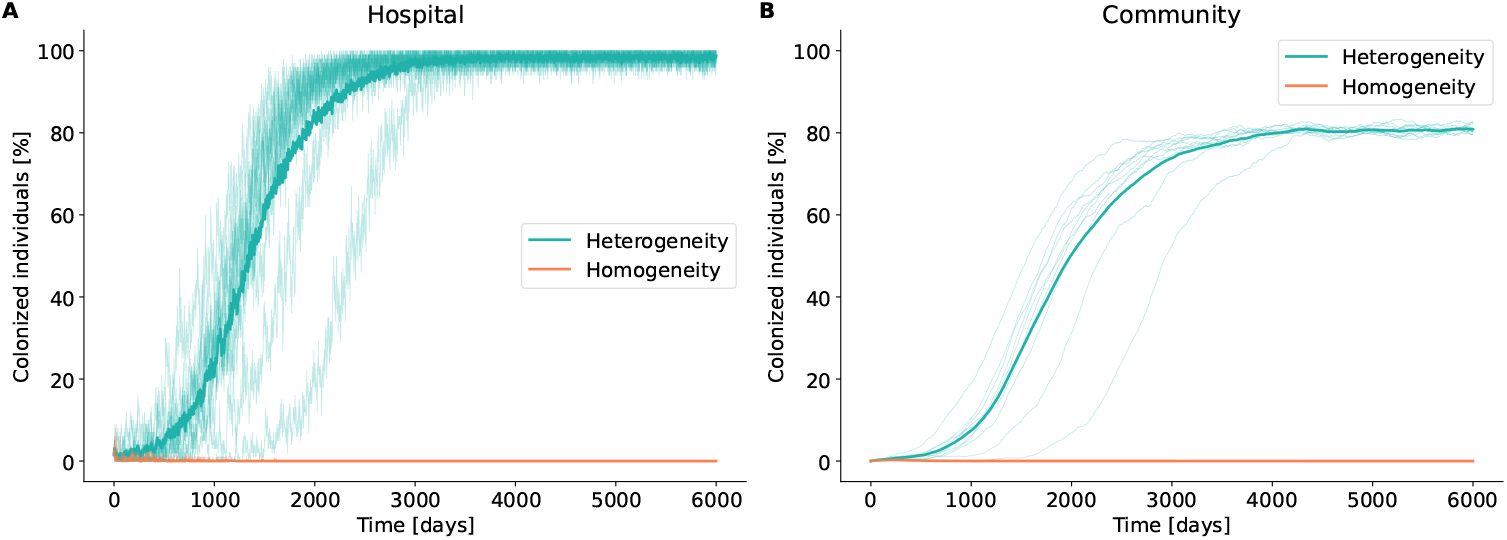
Colonization dynamics over time with and without population heterogeneity. Time series showing the percentage of individuals colonized by resistant bacteria under two conditions: with population heterogeneity and without. Darker shades represent average values, while lighter shades depict individual simulations (*n* = 10). Results are shown for the two settings: (A) hospital and (B) community.

### Detection sensitivity drives underestimation of resistance prevalence

The proportion of detected colonized individuals varies significantly depending on the sensitivity of the detection method employed. We compare two detection scenarios: an idealized screening with perfect sensitivity that detects any presence of resistant bacteria (*R* ≥1), and a more realistic approach requiring for detection a threshold of resistant bacteria (*R* ≥10^6^, detection limit estimated as described in the *Materials and Methods* section). Hereafter, we define individuals carrying *R* ≥10^6^ as *detectable* and those with 1 ≤ *R <* 10^6^ as *undetectable*. Notably, the choice of detection sensitivity substantially influenced prevalence estimates. In the hospital setting, 99% of individuals are colonized, yet only 60% are detectable (Fig. 3A). In the community the discrepancy is even more pronounced, with 81% of individuals colonized but only 10% detectable (Fig. 3D). These findings suggest that conventional detection techniques may substantially underestimate the true prevalence of individuals carrying resistant bacteria.

**Figure 3.**
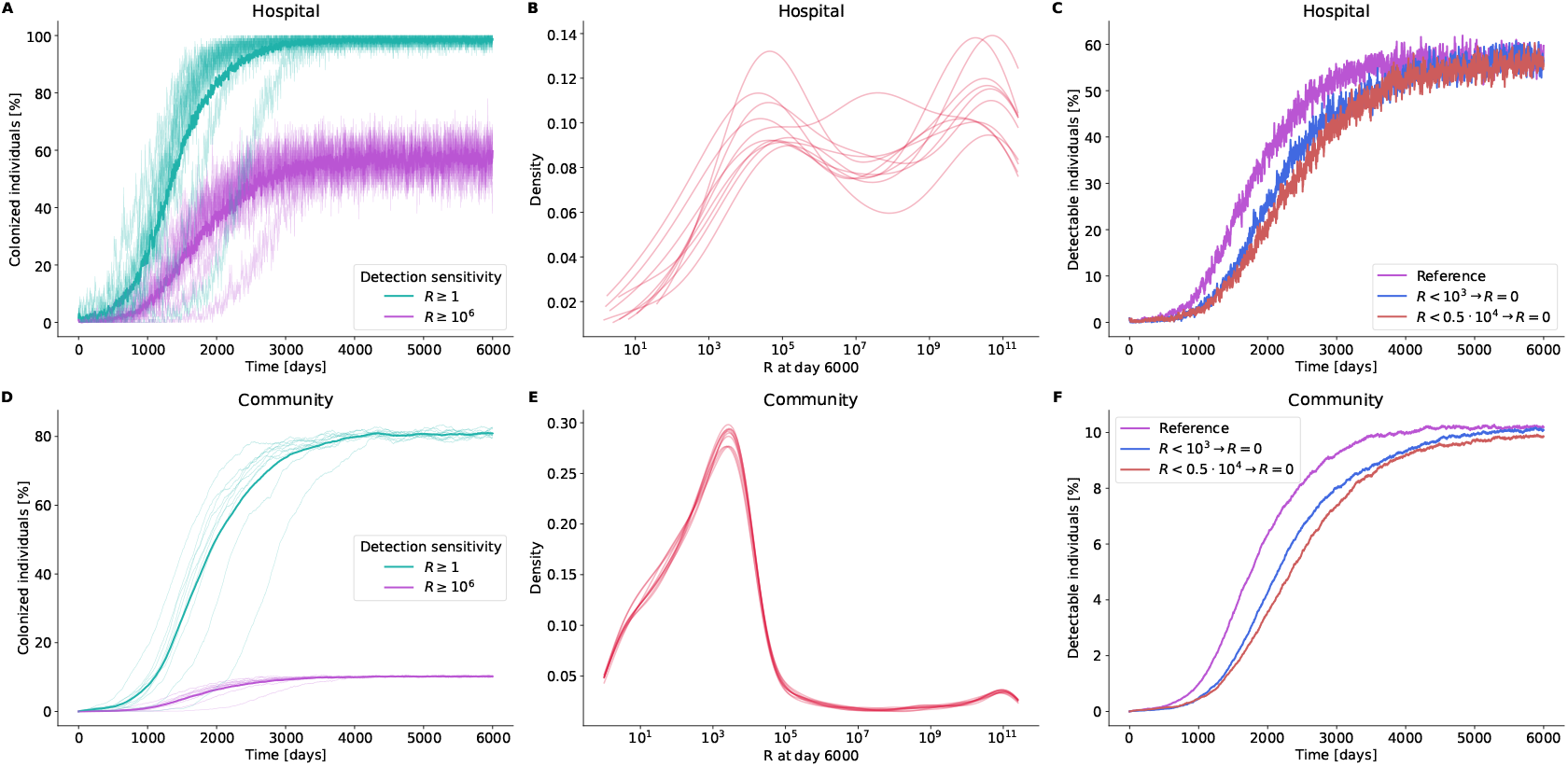
Detection sensitivity and the epidemiological role of undetectable carriers. (A, D) Time series showing the percentage of individuals colonized by resistant bacteria under two detection methods: an idealized screen detecting any resistant bacteria (*R* ≥1) and a realistic screen requiring a detection threshold (*R* ≥10^6^) in (A) hospital and (D) community. (B, E) Distributions of resistant bacteria among colonized individuals at day 6000 in (B) hospital and (E) community. (C,F) Time series of detectable individuals in hospital (C) and community (F) under the reference condition and scenarios in which individuals below 10^3^ or 0.5 × 10^4^ resistant bacteria are set to zero. For clarity, only average trajectories are shown. Darker shades denote means, lighter shades show individual simulations (*n* = 10).

#### Distribution of resistance load highlights detection bias

The final distribution of the resistance load (*R*) among colonized individuals provides insight into the impact of the detection threshold. Notably, in both hospital and community settings, resistance loads are concentrated near the transmission inoculum size (i.e. 10^4^), suggesting that many individuals carry resistant bacteria primarily due to recent transmission events (Fig. 3B,E). Lowering the detection threshold by at least two orders of magnitude (i.e. 10^6^ → 10^4^) would substantially increase the number of detected individuals in both environments.

#### Undetectable individuals delay resistance progression

To address the potential underestimation of colonized individuals, we investigate the influence of undetectable individuals on epidemiological dynamics. In these simulations, resistant bacteria counts below specific thresholds are set to zero to assess their impact. In both hospital and community settings, reducing the resistance load of low-level resistance carriage individuals results in slower epidemiological progression, while the final plateau levels are only slightly and mostly insignificantly lowered (Fig. 3C,F and Fig. S2). In the hospital, sigmoidal fits of colonization trajectories show that the midpoint is delayed by 20.9% and 30.6% under the two conditions, while plateau values decrease by 2.0% and 4.8%, respectively (Fig. S2C and Table S1). In the community, the midpoint is delayed by 21.4% and 32.4%, whereas plateau levels decline by 0.8% and 3.3%, respectively (Fig. S2D and Table S1). The results reveal a discernible epidemiological effect on the initial dynamic of resistance when undetectable individuals are disregarded, suggesting that even undetectable levels of colonization can contribute to the transmission of resistance. However, their influence on the persistence phase of resistance appears limited, This highlights the potential role of undetectable individuals in shaping the overall dynamics of resistance spread.

### Within-host coexistence of sensitive and resistant bacteria

The cross-scale nature of the model allows to capture the potential of within-host coexistence of resistant and sensitive bacteria. At the end of the simulation, we observe that among both colonized and detectable individuals, there is a wide range of relative frequency of resistance (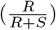) in both hospital and community settings (Fig. 4). The hospital and community populations do not segregate into distinct clusters of individuals exclusively colonized by resistant or sensitive bacteria. Instead, individuals are distributed across a continuum of relative resistance frequencies, with a higher density observed near low frequencies, suggesting that many individuals harbor only a small number of resistant bacteria compared to the sensitive community. This model thus captures the potential for coexistence of resistant and sensitive bacteria both within and between individual, reflecting a complex spectrum of colonization rather than discrete categories.

**Figure 4.**
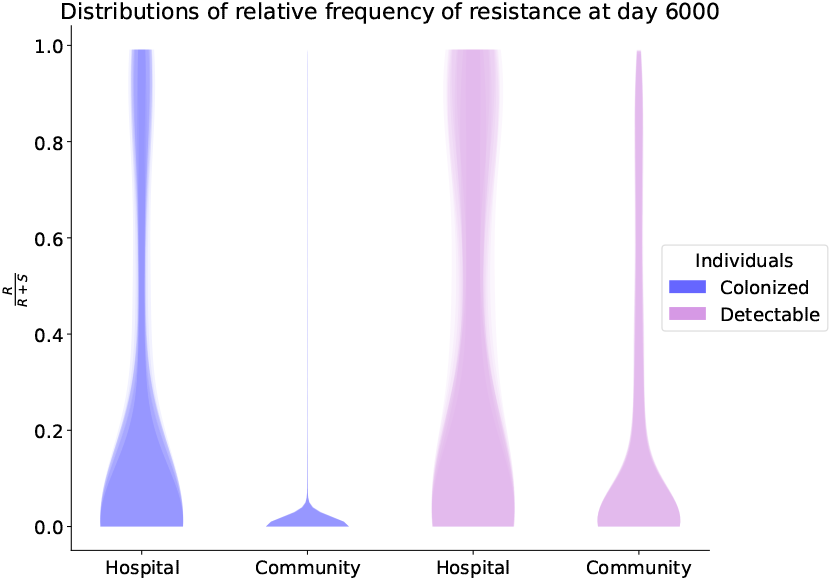
Relative frequency of resistance among colonized and detectable carriers at simulation end. Violin plots show the distribution of the within-host relative frequency of resistance 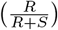 at day 6000 for both settings (hospital and community), displayed separately for colonized and detectable individuals. Overlaid violins correspond to independent simulation replicates (*n* = 10).

### Evaluating prevention measures

Strategies to prevent AMR range from antibiotic stewardship to interventions aimed at limiting transmission and reducing colonization. Here, we evaluate the impact of two hospital-based measures designed to reduce resistant bacteria: (i) isolation of resistant patients and (ii) intestinal decolonization.

#### Effectiveness of hospital isolation depends on transmission dynamics

We evaluate the impact of hospital entry screening and isolation on the system’s epidemiological dynamics. The intervention involves screening each new patient upon admission. Patients with resistant bacteria exceeding 10^7^ were isolated, preventing them from participating in transmission events as either donors or recipients. The threshold of 10^7^ resistant bacteria is chosen to ensure that resistance could be reliably detected during screening. Isolated patients are screened daily, and isolation is discontinued upon a negative screening result (*R* ≤ 10^7^). Comparative analysis of scenarios with and without this intervention demonstrated a substantial effect on epidemiological dynamics in both hospital and community settings (Fig. 5A,B). The magnitude of the intervention’s impact depended on the transmission probability within the hospital and the community. Simulations across varying transmission rates demonstrate the most pronounced effect when hospital transmission is high and community transmission is low, consistent with expectations, as isolation is most effective where transmission predominates [41]. In contrast, when community transmission rates increase, the influence of this hospital-based intervention diminishes drastically. Overall, these findings highlight the importance of isolating colonized patients in hospitals to control antibiotic resistance, particularly in high-transmission settings, while highlighting the interplay between hospital and community dynamics.

**Figure 5.**
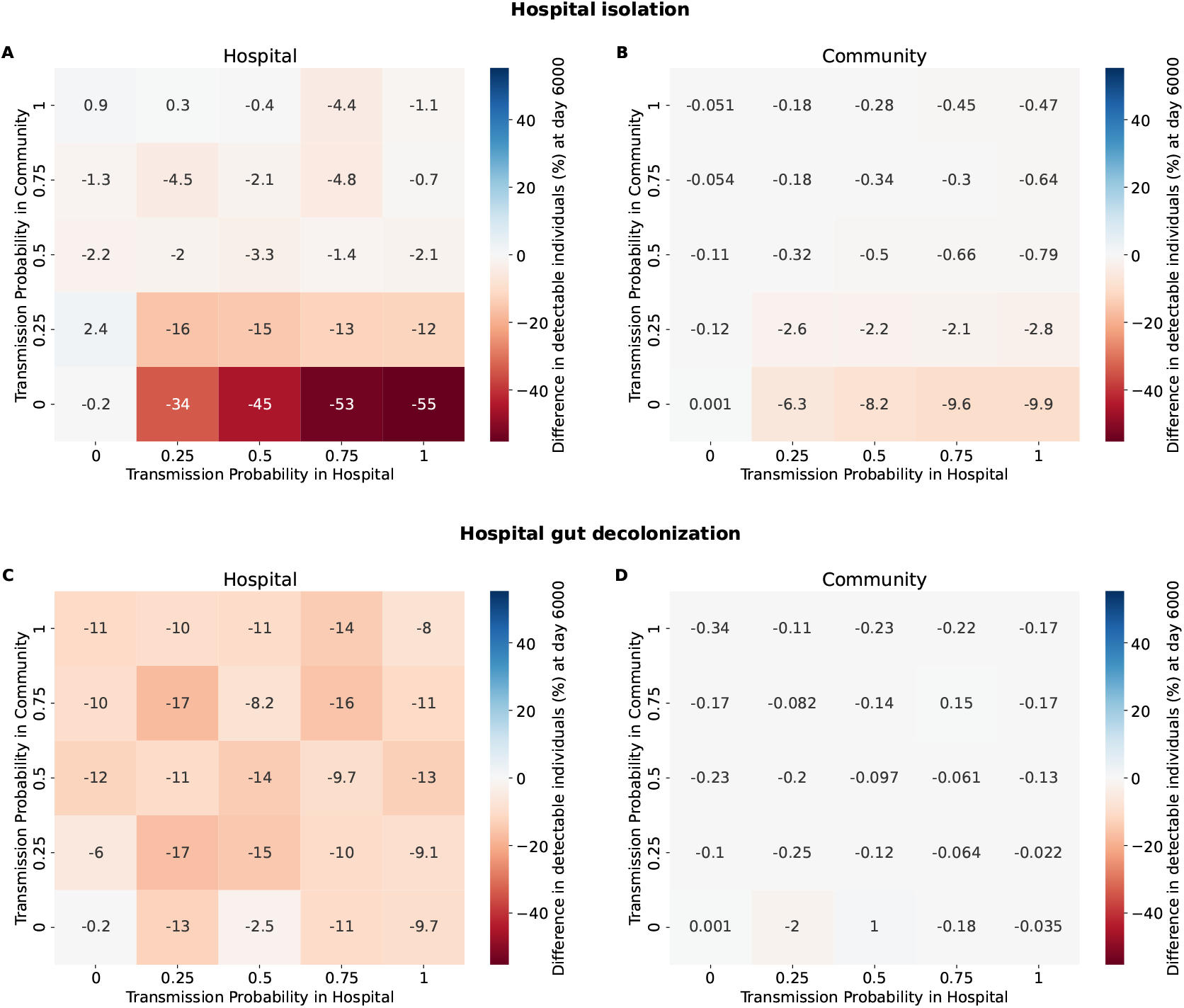
Impact of hospital isolation and gut decolonization on resistance colonization. Panels show the difference in colonization percentages (intervention minus reference) among detectable carriers (*R* ≥10^6^) at simulation end (day 6000), averaged across *n* = 10 simulations for each hospital–community transmission combination. (A,B) Patient isolation in hospital and community settings. (C,D) Hospital gut decolonization (1000-fold reduction of resistant bacteria per treatment) in hospital and community, respectively.

#### Hospital gut decolonization lowers prevalence independent of transmission dynamics

Multiple strategies have been proposed to reduce intestinal colonization by resistant bacteria, including: (i) selective decontamination of the digestive tract (SDD) using targeted antibiotics, (ii) the use of pre and probiotics, fecal microbiota transplantation, (iv) the use of specific phages, and (v) engineered CRISPR-Cas systems [42]. The choice of intervention depends on the specific bacteria and resistance mechanism involved. Here, we focus on SDD as a decolonization strategy targeting ESBL-producing bacteria. Data from meta-analyses and clinical trials show that SDD can temporarily reduce rectal carriage of ESBL-producing Enterobacterales, but sustained decolonization is uncommon and recolonization rates are high after cessation of therapy [43]. In our model, patients are screened at hospital admission, and those with resistant bacterial counts exceeding 10^7^ were selected for SDD. The intervention is implemented as a one-day reduction of both resistant and sensitive bacterial by 1000-fold. Simulations reveal that SDD significantly reduces hospital prevalence, with minimal impact on community resistance (Fig. 5C,D). In the hospital, the effectiveness of the intervention is consistent across different transmission rates, suggesting that SDD’s impact is largely independent of transmission intensity (Fig. 5C). Additional simulations with a milder decolonization of 100-fold reduction, resulting in quantitatively smaller yet qualitatively similar trends (Fig. S3).

### Sensitivity analysis

We evaluate the influence of four key parameters on colonization prevalence at simulation end: resistance cost, hospital treatment probability, community treatment rate, and inoculum size. Parameter importance was quantified using Sobol’ indices, where the first-order index captures the main effect of a parameter, and the total-order index additionally accounts for higher-order interactions. Full parameter ranges and details of the sensitivity analysis are provided in *Materials and Methods*. In the hospital setting, first- and total-order indices indicate that treatment rates and inoculum size exert the strongest main effect, while resistance cost contributed to a lesser extent (Fig 6A). In the community, inoculum size emerges as the dominant parameter, accounting for the largest share of output variance in both first- and total-order indices (Fig 6B). The effect of individual parameters on colonization prevalence, averaged over the remaining three parameters, reveals contrasting trends (Fig. S4). Increasing the resistance cost reduces the proportion of colonized individuals (Fig. S4A), whereas higher hospital treatment probability, community antibiotic consumption rate, and inoculum size all promote colonization in both compartments (Fig. S4B, S4C, S4D).

**Figure 6.**
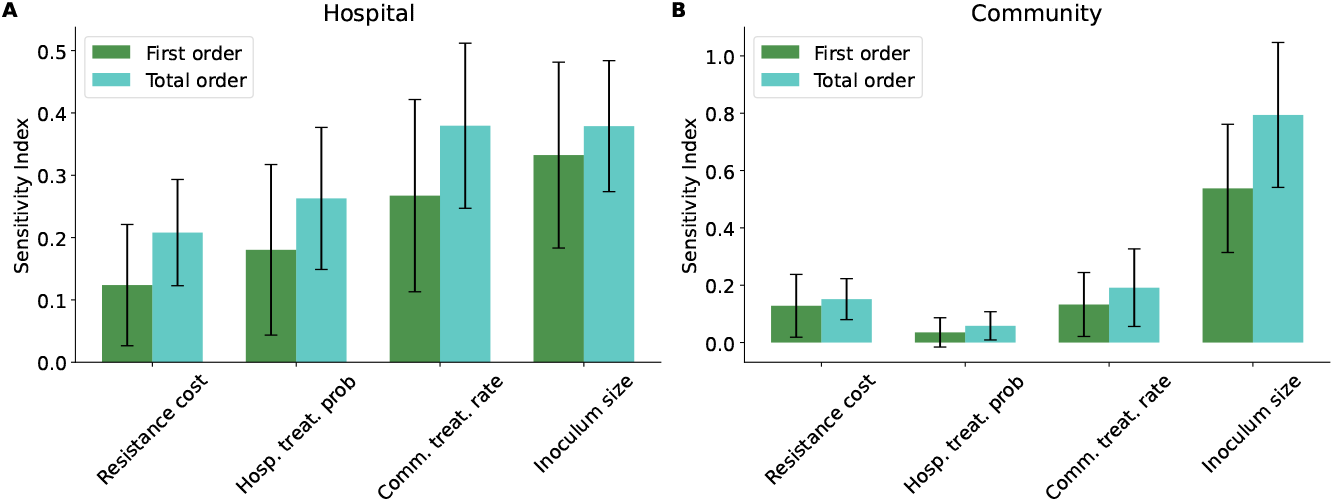
Sobol’ sensitivity analysis of resistance dynamics across settings. Global sensitivity indices quantifying the influence of key parameters on the proportion of detectable colonized individuals at simulation end (day 6000). (A) Hospital and (B) community settings showing first-order and total-order Sobol’ indices for resistance cost, hospital treatment probability, community treatment rate, and inoculum size. Error bars denote 95% confidence intervals across replicates (*n* = 10).

## Discussion

This study presents a cross-scale model of antimicrobial resistance that integrates within-host microbiota dynamics with population-level transmission in connected hospital and community settings. Although parameterized here to investigate the dissemination of extended-spectrum beta-lactamase (ESBL)-producing bacteria, the framework is adaptable to other resistance mechanisms, or even to multiple co-occurring resistances.

Our results highlight population heterogeneity as a key determinant of long-term resistance persistence across connected hospital and community settings. Even in scenarios with low but non-zero transmission, heterogeneity in antibiotic exposure and healthcare admissions was sufficient to maintain non-zero prevalence levels, while homogeneous populations fail to sustain resistance. This result is consistent with previous models that have identified population heterogeneity as a key factor in the persistence of antibiotic resistance [44–48]. This suggests that the same total amount of antibiotic use can result in vastly different scenarios depending on consumption patterns within the population. More broadly, these findings underscore the importance of incorporating individual-level variability into AMR models, since neglecting heterogeneity risks underestimating both persistence potential and the difficulty of eradication.

Our simulations demonstrate that detection thresholds can profoundly bias prevalence estimates, with realistic screening sensitivity underestimating colonization by up to an order of magnitude in the community. This gap arises in our simulations because a substantial fraction of carriers harbors resistant populations below the detection threshold and thus would remain undetectable in standard surveillance. These individuals, although below detection limits, can still contribute to the overall epidemiology. While their impact on long-term persistence appears limited in our model, their role in sustaining transmission chains suggests that current surveillance strategies may substantially underestimate the risk of spread. This hidden reservoir may limit the effectiveness of interventions and bias surveillance metrics, underscoring the need for diagnostics with lower detection thresholds.

Our results reveal that both hospital and community populations exhibit a continuum of within-host resistance frequencies, reflecting a complex spectrum of colonization rather than discrete categories. This approach differs from the classic compartmental models (e.g., SI frameworks), where the system allows for only one type of colonization [49] and aligns with empirical observations of mixed-strain colonization in the gut [50]. By explicitly coupling within-host bacterial dynamics to population-level transmission, our cross-scale framework captures the potential for the coexistence of resistant and sensitive bacteria, offering a mechanistic explanation for the persistence of resistance.

This modeling framework provides a tool to explore the potential impact of intervention strategies that would otherwise be difficult or expensive to test in real-world settings. By explicitly linking within-host colonization dynamics to population-level transmission, the model allows systematic evaluation of how interventions interact with hospital and community epidemiology. Our simulations highlight that intervention outcomes are strongly shaped by the context in which they are applied. Hospital isolation is predicted to be highly effective in reducing resistance when hospital transmission is dominant, yet its influence diminishes once community transmission became substantial. This underscores that hospital-based measures alone are insufficient if there is frequent transmission in the community, reinforcing the need for coordinated approaches across settings. By contrast, intestinal decolonization has a consistent effect on hospital prevalence regardless of transmission intensity. This suggests that interventions targeting carriage within individuals may be less dependent on external epidemiological conditions. However, the simulated reductions do not substantially alter community prevalence, highlighting the limitations of hospital reduction strategies in achieving community clearance.

The sensitivity analysis reveals clear differences in the determinants of resistance dynamics between hospital and community settings. In the hospital, both treatment rates and inoculum size exerted the strongest influence on resistance prevalence, underscoring that antibiotic exposure and transmission intensity jointly modulate hospital resistance levels. This finding highlights that hospital dynamics are not only governed by local treatment practices but also by the inflow of resistant bacteria selected in the community. In the community, the inoculum size emerged as the dominant driver of resistance prevalence, suggesting that transmission rather than selective pressure shapes long-term resistance persistence in low-treatment environments. Together, these results emphasize that resistance dynamics are shaped by different processes across scales: while hospital prevalence is primarily governed by treatment and quantity of bacteria transmitted, community persistence depends more strongly on transmission intensity.

Despite its utility, the model has several limitations. First, the hybrid approach combining deterministic within-host dynamics with stochastic individual-based population processes comes at a high computational cost, particularly when tracking absolute bacteria counts over extended simulation periods. Second, the parametrization of such a cross-scale model requires detailed datasets spanning both microbiota composition and resistance dynamics in clinical and community settings. These data are often sparse, fragmented, or inconsistent across studies, necessitating assumptions and extrapolations that may not fully capture biological and epidemiological complexity. Third, the simplification of the gut microbiota to a single phylum of interest, although motivated by the focus on ESBL-producing bacteria, overlooks the broader ecological context in which resistance evolves. Other phyla, such as Firmicutes or Bacteroidetes, may reduce or amplify resistance dynamics through competition, metabolic interactions, or horizontal gene transfer [51, 52]. Excluding them may therefore lead to a wrong estimation of colonization prevalence. Moreover, the assumption that ESBL plasmids are transferable within Proteobacteria neglects the substantial host specificity observed in plasmid-host compatibility [53]. Restricting the range of potential recipients potentially reduces the rate of resistance dissemination compared to our model assumptions. Fourth, the implementation of transmission events assumes that only a single bacterial type is transmitted at a fixed inoculum size. This simplification likely underestimates the heterogeneity and stochasticity of real-world transmission. In the absence of detailed empirical data on inoculum variability and composition during transmission, these assumptions remain necessary, but introduce uncertainty. Fifth, in practice gut decolonization does not uniformly reduce both sensitive and resistant Proteobacteria. However, in our model we assume that an effective decolonization strategy would affect them equally.

The key strength of this model is its explicit integration of within-host microbiota dynamics with human population transmission processes. Despite the limitations mentioned above, the model provides a clinically parameterized framework to explore the interconnected dynamics of the human microbiota and AMR epidemiology. Together, our findings show that resistance persistence is driven by the interplay of within-host dynamics, population heterogeneity, and setting-specific selective pressures. By linking individual colonization to hospital and community transmission, the framework provides a powerful tool for rapid and targeted testing of interventions under realistic epidemiological conditions.

## Materials and Methods

### Cross-scale gut microbiota model

Understanding the spread of antibiotic resistance requires integrating bacterial dynamics at multiple biological scales. We develop a cross-scale model linking within-host bacterial populations to population-level transmission in hospital and community environments. Below, we describe the within-host dynamics, the structure of the hospital and community populations, and the transmission mechanisms modeled.

#### Within-host gut microbiota

We model the within host dynamic of sensitive (S) and resistant (R) bacteria of the Proteobacteria phylum using ordinary differential equations (ODEs), according to the previosly develop ODE gut microbiota model developed by Tepekule et al [28] (Eq. 1 and 2). This focus on Proteobacteria is motivated by their high prevalence of clinically significant antibiotic resistance in species such as *Escherichia coli, Klebsiella pneumoniae, Pseudomonas aeruginosa*, and *Enterobacter* species [35]. By restricting our model to Proteobacteria, we balance biological realism with model tractability. The model incorporates growth, interaction, plasmid conjugation, plasmid segregation and antibiotic effect. Bacterial growth of the sensitive bacterial population follows mass-action kinetics at rate *µ*_*S*_. The growth of the resistance population (*µ*_*R*_) is reduced compared to the one of the sensitive one by a fitness cost (*ρ*) imposed by resistance carriage, such that: *µ*_*R*_ = *µ*_*S*_(1 − *ρ*). The interaction rate, *a*, accounts for the effect of overall gut bacterial density on population growth. Plasmid segregation occurs at rate *γ*, representing the fraction of resistant bacteria that lose the plasmid during replication. Plasmid conjugation, denoted by rate *h*, facilitates horizontal transfer of resistance genes. Antibiotic effect, applied only during treatment, selectively reduces the sensitive bacterial population with death rate *δ*. The resulting dynamics are described by the following system of differential equations:

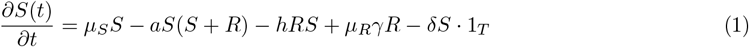

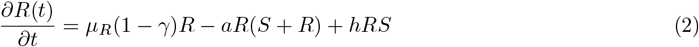

Here, *S*(*t*) and *R*(*t*) represent the sensitive and resistant bacterial populations at time *t*, and 1_*T*_ is an indicator function that equals 1 during antibiotic treatment and 0 otherwise. In the absence of antibiotics, the gut sensitive population converges to the equilibrium 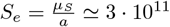 bacteria.

#### Community and hospital environments

To construct the cross-scale model, we consider a population composed of multiple individuals whose microbiota follows Eq. 1 and 2 (Fig. 1). To account for the differences in antibiotic consumption patterns and transmission dynamics across settings, we introduce two distinct environments: the hospital and the community. The hospital environment has a fixed population size of *N*_*h*_ = 10^2^ individuals, while the community comprises *N*_*c*_ = 10^4^ individuals. Hospitalized patients are assigned a specific duration of stay and an antibiotic consumption profile upon admission. To maintain stable population sizes in both environments, discharged hospital patients are replaced by randomly selected individuals from the community, thereby mimicking the constant flux between hospital and community populations.

#### Stochastic transmission events

Bacterial transmission within each environment (hospital or community) occurs stochastically, with transmission rates specific to each setting. Transmission probabilities per step are set at *p*_*c*−*trans*_ = 0.25 for the community and *p*_*h*−*trans*_ = 1 for the hospital, reflecting the higher risk of transmission in hospital settings due to closer patient contact and increased exposure to contaminated surfaces. When a transmission event is triggered, the donor transfers a fixed amount of 10^4^ bacterial bacteria to a randomly chosen recipient within the same environment (Fig. 1). The composition of the transferred bacteria (all sensitive or all resistant) depends on the bacterial composition of the donor. Specifically, the relative frequency of resistant bacteria in the donor’s gut microbiota determines the likelihood that the transferred bacteria will be resistant. This frequency is calculated as 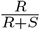, where *R* and *S* denote the resistant and sensitive bacterial populations in the donor at the time of transmission. This transmission process abstracts real-world mechanisms and does not differentiate between direct (e.g., person-to-person) and indirect (e.g., via surfaces) transmission routes. Instead, it captures the overall probability and scale of bacterial dissemination within each environment.

#### System update

Simulations are initiated with a single hospital patient highly colonized by resistant bacteria (*R* = 10^11^, *S* = 0), while all other individuals carry only sensitive bacteria (*R* = 0, *S* = 10^11^). Our model updates its state daily by sequentially executing the following processes:

- The bacterial counts within each individual are updated according to Euler’s approximation:

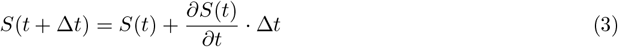

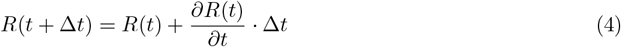

where the time step Δ*t* = 1 day.
- Untreated community members initiate outpatient antibiotic treatment at rate *p*_*c*−*treat*_.
- Hospital patients that complete their stay are discharged to the community and replaced by a randomly selected individual from the community.
- Newly admitted hospital patients are assigned to the treatment category with probability *p*_*h*−*treat*_ and given a corresponding duration of stay and, if relevant, an antibiotic treatment duration.
- Bacterial transmission events occur, following the stochastic process described above.

### Parameter Estimation

#### Within-host bacterial behavior

The bacterial dynamics in Eq. 1 and 2 are parameterized based on values reported by Tepekule *et al*. [28]. These parameters, summarized in Table S2, include bacterial growth rate *µ*_*S*_, resistance costs *ρ*, and interaction term *a*, conjugation rate *h*, missegregation rate *γ* and antibiotic-induced death rate *δ*. The antibiotic killing rate is increased by a factor 2.35 such that a 3-day treatment leads to 10 fold reduction of sensitive bacteria from steady-state equilibrium [54].

#### Hospital antibiotic treatment probability

To estimate the probability of receiving antibiotics during hospitalization (*p*_*h*−*treat*_), we examine the proportion of hospitalizations, where beta-lactam antibiotics of interest were administered. Age-stratified data from 2022 and 2023 for the University Hospital Zurich (USZ) indicates stable consumption ratios across most age groups, with the exception of children under age 10, likely due to higher birth rates without antibiotic prescriptions (Fig. S5A). To avoid bias, we exclude this age group and calculate a weighted average consumption ratio for the remaining patients, resulting in *p*_*h*−*treat*_ = 0.4570.

#### Hospitalization duration

To model the distribution of the hospitalization duration in the USZ dataset for 2022 and 2023, we fit a lognormal distribution. This distribution is particularly suitable for positively skewed data (Fig. S5B). The probability density function of the lognormal distribution is given by:

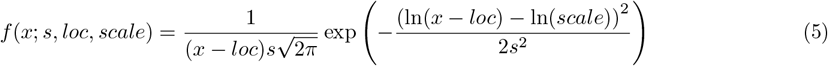

where *s* is the shape parameter, *loc* is the location parameter, and *scale* is the scale parameter. We fit separate lognormal distributions for patients who did and did not receive antibiotics, with the resulting parameter estimates shown in Table S2.

#### Hospital antibiotic treatment duration

To determine the total antibiotic duration of each patient in the antibiotic group, we extract the total number of days receiving antibiotics based on the distribution derived from their hospitalization duration. This is inferred from the distribution of treatment durations, derived from the USZ surveillance dataset for the years 2022 and 2023 (Fig. S5C).

#### Community consumption

Outpatient antibiotic consumption rates are obtained from the Swiss Center for Antibiotic Resistance (ANRESIS) database [39]. We focus on beta-lactam antibiotics, to which ESBL-producing bacteria are resistant (Table S3). The most recent available data (2023) indicates an overall consumption rate of 4.77 daily defined doses (DDD) per 1,000 inhabitants per day. For simplicity, we assume that 1 DDD corresponds to one person under treatment. Based on typical treatment patterns, we further estimate an average duration of antibiotic therapy of 5 days [55, 56]. At each update an untreated individual in the community may start the antibiotic treatment with fixed rate *p*_*c*−*treat*_. In order to reflect the average of 4.77 ‰ individuals under treatment every day, knowing that the treatment duration is 5 days and that at any time (even during treatment) an individual may go to the hospital we need to compute the effective outpatient treatment duration (*d*_*treat*_). The probability of going to the hospital (*p*_*c*→*h*_) is dependent on the average hospital stay (*µ*_*h*_) and environments sizes (*N*_*h*_ and *N*_*c*_): 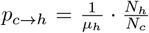. The effective outpatient treatment duration can be written as:

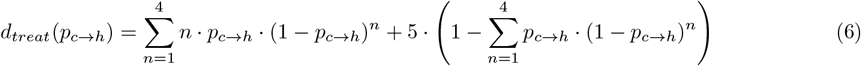

At each step, the individual treatment starting rate in the community becomes: *p*_*c*−*treat*_ = 4.77‰*/d*_*treat*_.

#### Population heterogeneity

To capture population-level variability in antibiotic consumption, we model outpatient antibiotic consumption heterogeneity based on Olesen *et al*., which showed that 10% of the population accounts for 60% of outpatient antibiotic use, while 24% consumes the remaining 40% [36]. We therefore create 3 categories of individuals and assign each person to one of them according to the previously defined distribution. At simulation start, the highly colonized individual is assigned to the top 10% of the population responsible for 60% of antibiotic consumption. We further incorporate non-random community-hospital transitions, meaning that upon discharge of a hospital patient the replacement is not picked randomly from the community environment but depending on which of the three categories the individuals are in the chance of being picked is different. This reflects the increased likelihood of hospitalization for individuals with higher antibiotic consumption (e.g. more vulnerable groups). Consequently, 66% of the population neither visits the hospital nor uses antibiotics, simulating a lower-risk segment of the community. The state incorporating community consumption variability and non-random transitions is referred to as the heterogeneous model, while the absence of such variability defines the homogeneous model. Similar to the homogeneous case, for the heterogeneity case, we need to compute the new probability of community treatment rates in class 1, 2 and 3. These depend on the probability of going to the hospital and the effective community treatment duration for each class, final values are reported in Table S2.

### Realistic detection sensitivity

To estimate the order of magnitude of the minimum number of resistant bacteria required for detection in standard rectal surveillance screening, we reconstruct the sequence of dilution steps from gut content to plated material (Fig. S6). First, assuming a total of approximately 10^14^ bacterial cells in the human gut [9] and an average fecal concentration of 10^11^ bacteria per gram [57], only about 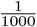 of the total gut population is excreted per gram of feces. Our empirical measurements further indicate that the rectal swab collects a median of ≃ 0.1 g of fecal material (Fig. S7). This material is then typically suspended in 1 *mL* of transport medium, and only 10 *µL* is subsequently plated for screening, corresponding to an additional 10^−2^ dilution. Hence, the total dilution from gut content to plated material is on the order of 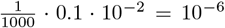. We further experimentally estimated that growth is detectable if at least one ESBL-producing bacterium (≃ *CFU* ) is present on the selective plate. The resulting detection limit corresponds to approximately 10^6^ resistant bacteria in the gut.

### Sensitivity analysis

We explore the effect of four key parameters on the percentage of detectable colonized individuals at the end of the simulation: the resistance cost, the hospital treatment probability upon admission, the daily community treatment rate, and the inoculum size transferred during each transmission event. The reference values and sampling ranges for these parameters are reported in Table 1. A Sobol’ sensitivity analysis is performed using 1280 parameter combinations generated with the Saltelli sampling scheme [58]. The analysis focuses on first-order and total-order Sobol’ indices of colonization prevalence in both hospital and community settings. The first-order measures the main effect of a single input variable on the output variance, while the total-order captures the total contribution of an input variable to the output variance, including both its main effect and all its interactions with other variables.

**Table 1:**
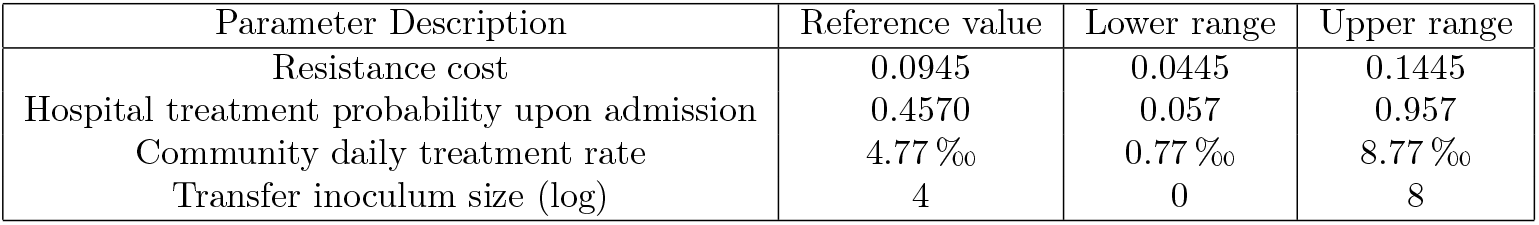
Parameter ranges used in the Sobol’ sensitivity analysis. Reference values and sampling ranges for the four parameters explored: resistance cost, hospital treatment probability, community treatment rate, and inoculum size.

| Parameter Description | Reference value | Lower range | Upper range |
| --- | --- | --- | --- |
| Resistance cost | 0.0945 | 0.0445 | 0.1445 |
| Hospital treatment probability upon admission | 0.4570 | 0.057 | 0.957 |
| Community daily treatment rate | 4.77 ‰ | 0.77 ‰ | 8.77 ‰ |
| Transfer inoculum size (log) | 4 | 0 | 8 |

## Code availability statement

All code used in this study will be made publicly available on GitHub upon publication. Prior to publication, the code can be obtained from the corresponding author upon request.

## Declaration of generative AI use

During the preparation of the manuscript, ChatGPT (GPT-4o, 5, 5.1) by OpenAI was employed for text editing purposes.

## Acknowledgments

We would like to thank Christopher Witzany and Lorenzo Talamanca for helpful discussions and comments.

## Notes

### Competing Interest Statement

The authors have declared no competing interest.

