## Supplementary Tables and Figures for "Cross-scale effect of microbiota dynamics on resistance"

### Supplementary material

#### S1 Supplementary tables

| Setting / Condition | $L$ | $k$ | $x_0$ | $b$ | $R^2$ |
|---|---|---|---|---|---|
| *Community* | | | | | |
| Reference | 10.655 | 0.002109 | 1757.4 | -0.524 | 0.9988 |
| $R < 10^3 \to R = 0$ | 10.439 | 0.001785 | 2125.4 | -0.540 | 0.9976 |
| $R < 0.5 \cdot 10^4 \to R = 0$ | 10.144 | 0.001723 | 2296.0 | -0.380 | 0.9992 |
| *Hospital* | | | | | |
| Reference | 59.575 | 0.002113 | 1705.8 | -2.972 | 0.9949 |
| $R < 10^3 \to R = 0$ | 59.111 | 0.001759 | 2071.7 | -3.371 | 0.9938 |
| $R < 0.5 \cdot 10^4 \to R = 0$ | 57.615 | 0.001715 | 2251.8 | -2.299 | 0.9957 |

Table S1: **Sigmoid fit parameters for colonization dynamics.** The fitted function is $f(x) = \frac{L}{1+\exp\left(-k(x-x_0)\right)} + b$, where $L$ is the maximum value, $k$ the steepness, $x_0$ the midpoint, $b$ the offset, and $R^2$ the coefficient of determination.

| Parameter | Description | Equation or Value |
|---|---|---|
| **Within-host bacterial behavior** | | |
| $\mu_S$ | Growth rate of sensitive bacteria | 0.27198 [1] |
| $\rho$ | Resistance cost | 0.0945 [1] |
| $\gamma$ | Missegregation fraction | 0.0227 [1] |
| h | Conjugation frequency | $10^{-14.8}$ [1] |
| a | Bacterial interaction | $\frac{9.3}{10^{13}}$ [1] |
| $\delta$ | Antibiotic death rate | $0.283 \cdot 2.35$ [1, 2] |
| **Hospital** | | |
| $N_h$ | Hospital size | $10^2$ |
| $p_{h-trans}$ | Probability of transmission event per day | 1 |
| $p_{h-treat}$ | Treatment probability upon admission | 0.4570 (Fig. S5A) |
| $s_t$ | Shape of the lognormal distribution for treated patients | 0.8571 (Fig. S5B) |
| $s_u$ | Shape of the lognormal distribution for untreated patients | 0.7893 (Fig. S5B) |
| $loc_t$ | Location of the lognormal distribution for treated patients | 0 (Fig. S5B) |
| $loc_u$ | Location of the lognormal distribution for untreated patients | 0 (Fig. S5B) |
| $scale_t$ | Scale of the lognormal distribution for treated patients | 5.8968 (Fig. S5B) |
| $scale_u$ | Scale of the lognormal distribution for untreated patients | 3.2448 (Fig. S5B) |
| $\mu_{h,t}$ | Average hospitalization duration for treated patients | $scale_t \cdot e^{\frac{s_t^2}{2}} + loc_t$ |
| $\mu_{h,u}$ | Average hospitalization duration for untreated patients | $scale_u \cdot e^{\frac{s_u^2}{2}} + loc_u$ |
| $\mu_h$ | Average hospitalization duration | $\mu_{h,t} \cdot p_{h-treat} + \mu_{h,u} \cdot (1 - p_{h-treat})$ |
| **Community** | | |
| $N_c$ | Community size | $10^4$ |
| $p_{c-trans}$ | Probability of transmission event per day | 0.25 |
| $\mu_t$ | Average treated individuals per day in the community | 4.77 ‰[3] |
| **Homogeneous case** | | |
| $p_{c\to h}$ | Probability of transitioning from community to hospital | $\frac{1}{\mu_h} \cdot \frac{N_h}{N_c}$ |
| $d_{treat}$ | Effective outpatient treatment duration | $\sum_{n=1}^{4} n \cdot p_{c\to h} \cdot (1 - p_{c\to h})^n + 5 \cdot \left(1 - \sum_{n=1}^{4} p_{c\to h} \cdot (1 - p_{c\to h})^n\right)$ |
| $p_{c-treat}$ | Treatment starting rate in the community | $\frac{\mu_t}{d_{treat}}$ . |
| **Heterogeneous case** | | |
| $f_1$ | Fraction of individuals in class 1 | 0.1 [4] |
| $f_2$ | Fraction of individuals in class 2 | 0.24 [4] |
| $f_3$ | Fraction of individuals in class 3 | 0.66 [4] |
| $a_1$ | Fraction of outpatient antibiotic consumed by class 1 | 0.6 [4] |
| $a_2$ | Fraction of outpatient antibiotic consumed by class 2 | 0.4 [4] |
| $p_{c_1\to h}$ | Transition probability for class 1 | $\frac{1}{\mu_h} \cdot \frac{a_1 \cdot N_h}{f_1 \cdot N_c}$ |
| $p_{c_2\to h}$ | Transition probability for class 2 | $\frac{1}{\mu_h} \cdot \frac{a_2 \cdot N_h}{f_2 \cdot N_c}$ |
| $p_{c_3\to h}$ | Transition probability for class 3 | 0 |
| $d_{treat,1}$ | Effective treatment duration for class 1 | $d_{treat}(p_{c_1\to h})$ |
| $d_{treat,2}$ | Effective treatment duration for class 2 | $d_{treat}(p_{c_2\to h})$ |
| $p_{c-treat1}$ | Treatment starting rate for class 1 | $\frac{1}{1000} \cdot \frac{a_1}{f_1 \cdot d_{treat,1}}$ |
| $p_{c-treat2}$ | Treatment starting rate for class 2 | $\frac{1}{1000} \cdot \frac{a_2}{f_2 \cdot d_{treat,2}}$ |
| $p_{c-treat3}$ | Treatment starting rate for class 3 | 0 |

Table S2: **Model parameters.** Overview of parameters used in the cross-scale model, including within-host bacterial dynamics, hospital and community compartments, and homogeneous and heterogeneous treatment scenarios. Values are based on published data where available, otherwise derived from fitted distributions or model assumptions.

| | Antibiotic | DDD ‰ inhabitants$^{-1}$ day$^{-1}$ |
|---|---|---|
| | 1st and 2nd generation Cephalosporins | 0.55 |
| | 3rd and 4th generation Cephalosporins | 0.07 |
| | Comb. of pen. (with anti-pseudomonal activity) | 0 |
| | Comb. of pen. (without anti-pseudomonal activity) | 2.85 |
| | Beta-lactamase resistant penicillins | 0 |
| | Beta-lactamase sensitive penicillins | 0.05 |
| | Penicillins with extended spectrum | 1.25 |
| Total | | 4.77 |

Table S3: **Outpatient antibiotic consumption in 2023 in Switzerland** [3]. Data are expressed as defined daily doses (DDD) per 1,000 inhabitants per day.

### S2 Supplementary figures

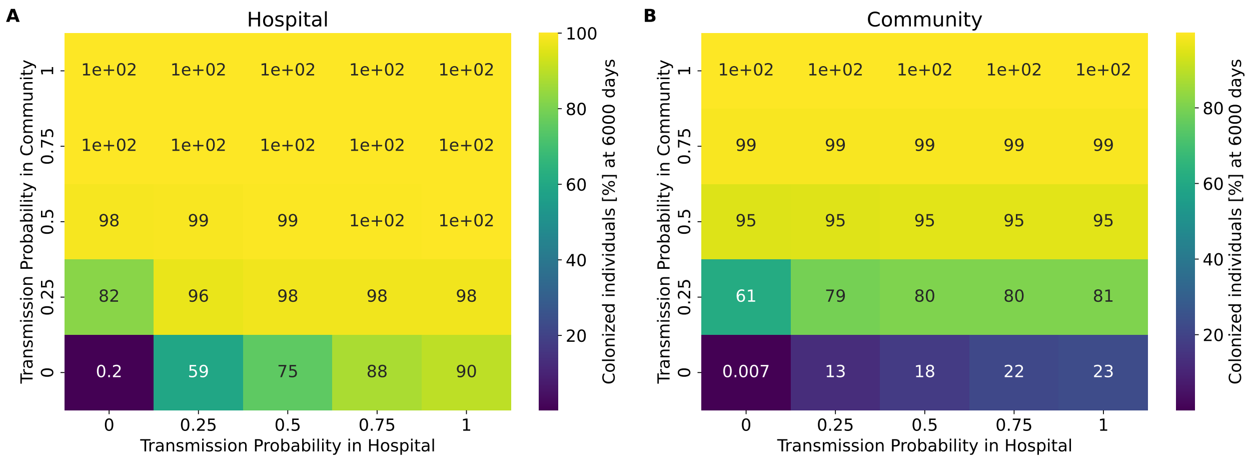


Figure S1: **Impact of transmission rates on colonization.** Colonization percentages in (A) hospital and (B) community settings, with different transmission rates and heterogeneity. For each pair of hospital and community transmission probabilities, the mean colonization rate is shown, averaged over 10 simulations at day 6000.

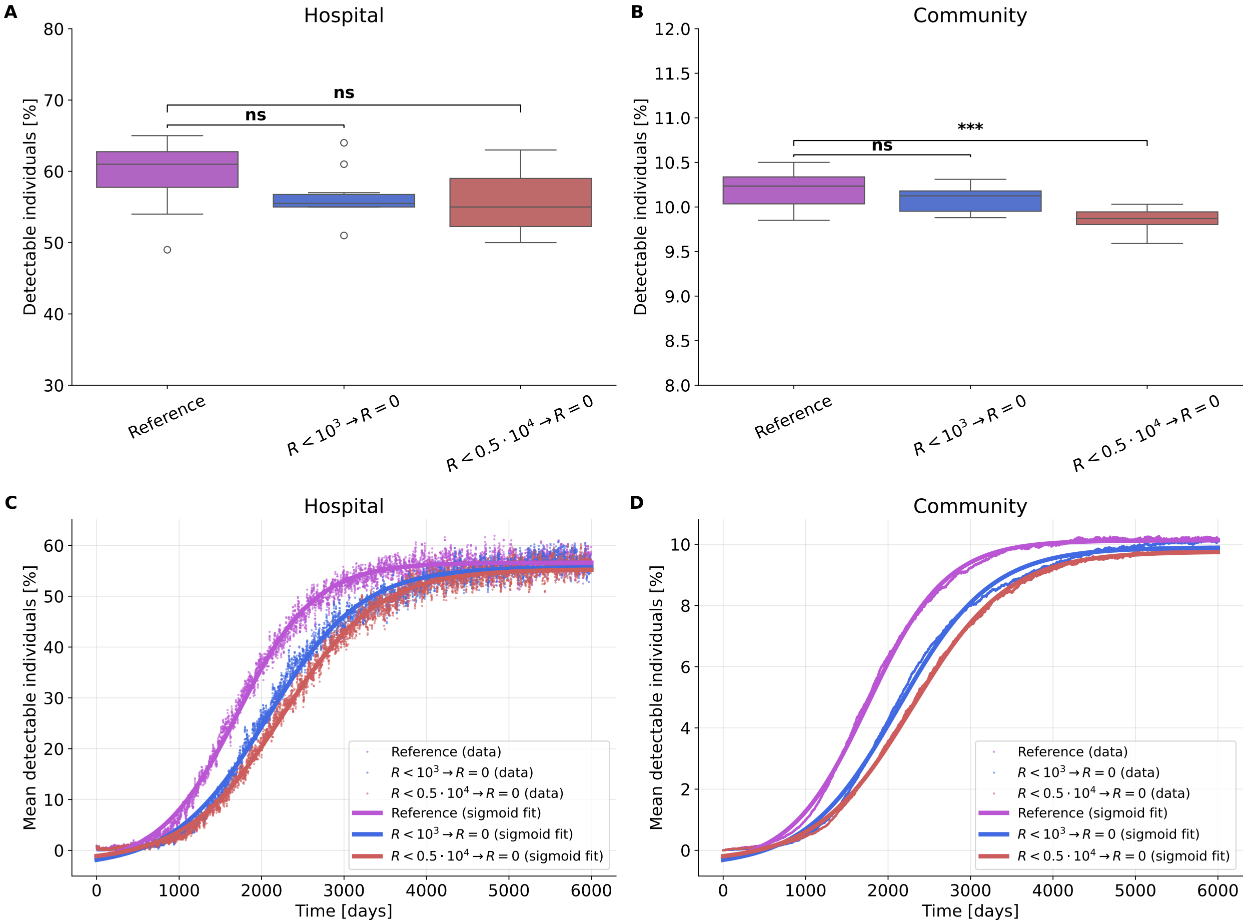


Figure S2: **Impact of *undetectable* individuals on colonization dynamics.** Three scenarios are compared: reference and two cases where individuals with resistant bacterial counts below $10^3$ or $0.5 \cdot 10^4$ were set to zero. (A, B) Final distribution (day 6000) of detectable individuals across 10 simulation replicates in (A) hospital and (B) community settings. Statistical significance was assessed using independent t-tests ($\alpha = 0.05$). Significance brackets indicate: *** $p < 0.001$, ** $p < 0.01$, * $p < 0.05$, ns = not significant. (C, D) Average trajectories of detectable individuals ($n = 10$) in (C) hospital and (D) community, with their sigmoidal fits. Fit parameters and goodness-of-fit ($R^2$) are reported in Table S1.

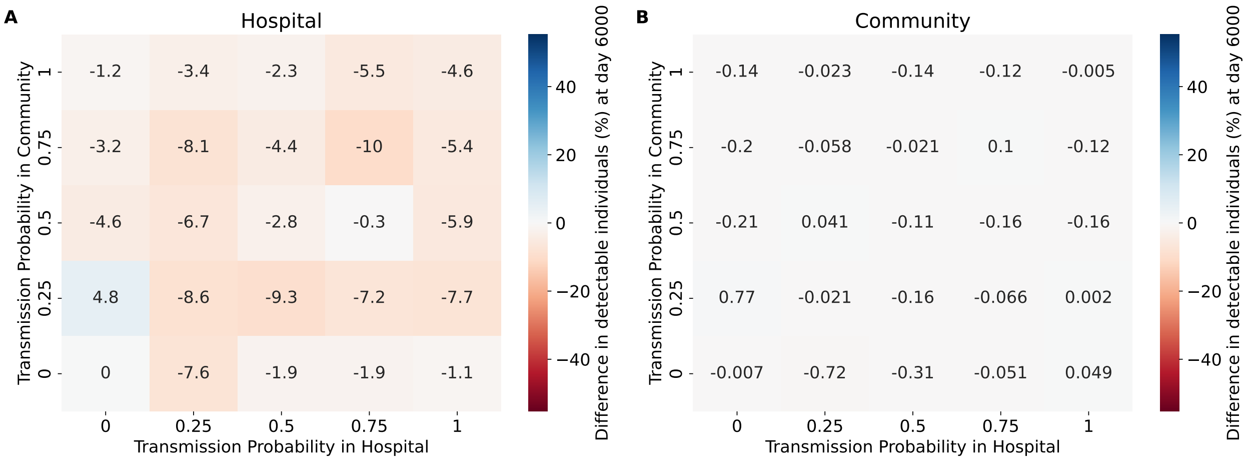


Figure S3: **Impact of hospital gut decolonization (100-fold) on colonization rates.** Difference in colonization percentages (intervention minus reference) among detected carriers ($R \geq 10^6$) at simulation end (day 6000), averaged across $n = 10$ simulations for each hospital–community transmission combination. (A–B) Gut decolonization (100-fold reduction of resistant bacteria per treatment) in hospital and community settings, respectively.

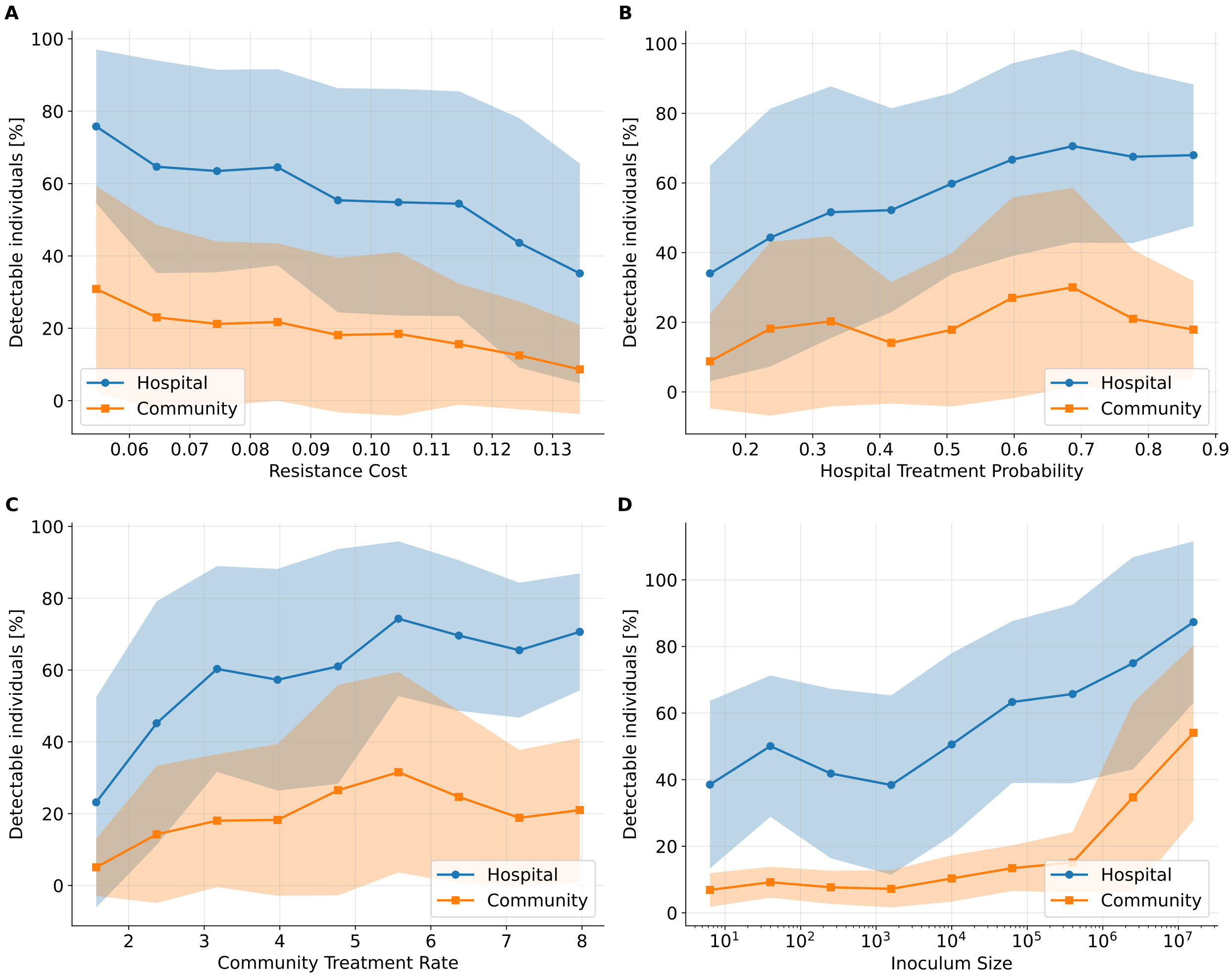


Figure S4: **Impact of sensitivity parameters on colonization prevalence**. Moving-window analysis of the proportion of colonized individuals ($R \geq 10^6$) at the end of the simulation (day 6000) as a function of (A) resistance cost, (B) hospital antibiotic treatment probability, (D) community antibiotic consumption rate, and (D) inoculum size. Solid lines represent mean values in hospital (blue) and community (orange) populations, with shaded bands indicating $\pm 1$ standard deviation. Each curve summarizes average results from 10 simulation replicates per parameter set (1280), smoothed using a rolling window spanning 20% of the parameter range.

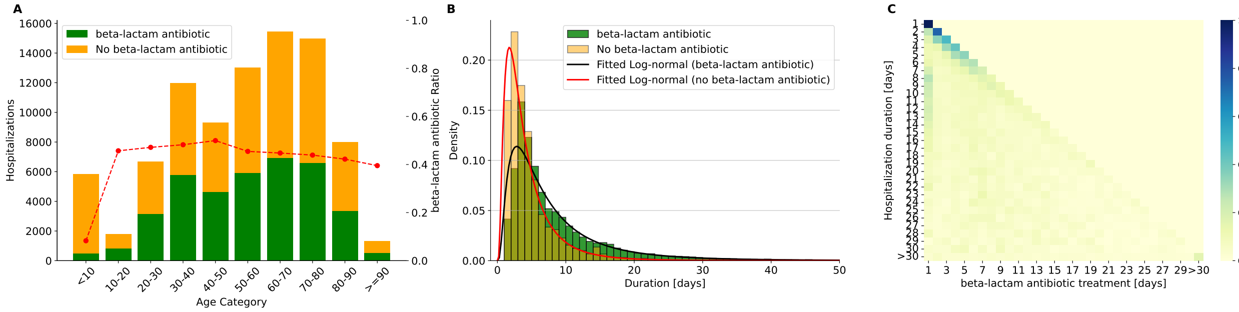


Figure S5: **Hospital patterns at the University Hospital Zurich (USZ) in 2022–2023.** (A) Age-based distribution of hospitalizations with consumption of ESBL-relevant antibiotics. Hospitalized patients are grouped into 10-year age categories. Yellow bars represent the total number of hospitalizations in each age group, green bars indicate hospitalizations with consumption of at least one ESBL-relevant antibiotic, and red dots show the proportion of hospitalizations with antibiotic consumption. (B) Distribution of hospitalization duration for patients treated (green) or not treated (yellow) with ESBL-relevant antibiotics. Fitted lognormal distributions are shown in red (treated) and black (untreated). (C) Heatmap showing the distribution of treatment days among treated patients, stratified by hospital stay duration.

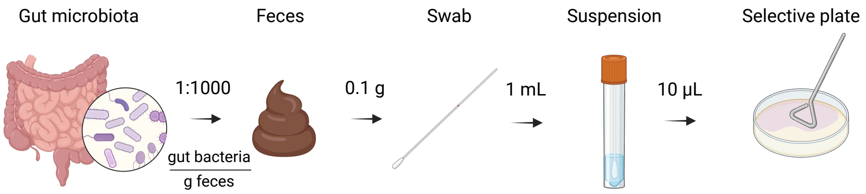


Figure S6: **Schematic representation of dilution steps from gut content to selective plate.** Only $\frac{1}{1000}$ of the gut bacterial population is excreted per gram of feces, of which a median of 0.1 g is collected by a rectal swab. The swab is suspended in 1 mL of transport medium, and 10 $\mu$L is plated for screening. Created in BioRender.

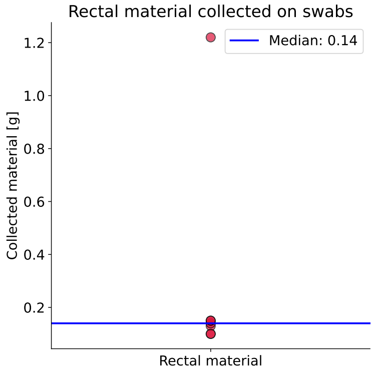


Figure S7: Weight of rectal material collected on surveillance swabs from patients at the University Hospital Zurich ($n = 7$). The median amount of collected material was approximately 0.14g, corresponding to an order of magnitude of 0.1g.
